# BOXR1030, an anti-GPC3 CAR with exogenous GOT2 expression, shows enhanced T cell metabolism and improved antitumor activity

**DOI:** 10.1101/2021.11.17.469041

**Authors:** Taylor L. Hickman, Eugene Choi, Kathleen R. Whiteman, Sujatha Muralidharan, Tapasya Pai, Tyler Johnson, Avani Parikh, Taylor Friedman, Madaline Gilbert, Binzhang Shen, Luke Barron, Kathleen E. McGinness, Seth A. Ettenberg, Greg T. Motz, Glen J. Weiss, Amy Jensen-Smith

**Affiliations:** Employees of Unum Therapeutics, Inc. Cambridge, MA USA while work was performed; Current SOTIO Biotech Inc. employees

**Author notes:** Corresponding authors: Glen J. Weiss, 200 Cambridge Park Dr. Cambridge, MA USA, Amy Jensen-Smith, 200 Cambridge Park Dr. Cambridge, MA USA. Equal contribution (Taylor L. Hickman, Eugene Choi).

**Keywords:** Glypican-3, immunometabolism, chimeric antigen receptor, T cell exhaustion, solid tumor

## Abstract

**Purpose:** The solid tumor microenvironment (TME) drives T cell dysfunction and inhibits the effectiveness of immunotherapies such as chimeric antigen receptor-based T cell (CAR T) cells. Early data has shown that modulation of T cell metabolism can improve intratumoral T cell function in preclinical models.

**Experimental Design:** We evaluated GPC3 expression in human normal and tumor tissue specimens. We developed and evaluated BOXR1030, a novel CAR T therapeutic co-expressing glypican-3 (GPC3)-targeted CAR and exogenous glutamic-oxaloacetic transaminase 2 (GOT2) in terms of CAR T cell function both *in vitro* and *in vivo*.

**Results:** Expression of tumor antigen GPC3 was observed by immunohistochemical staining in tumor biopsies from hepatocellular carcinoma, liposarcoma, squamous lung cancer, and Merkel cell carcinoma patients. Compared to control GPC3 CAR alone, BOXR1030 (GPC3-targeted CAR T cell that co-expressed GOT2) demonstrated superior *in vivo* efficacy in aggressive solid tumor xenograft models, and showed favorable attributes *in vitro* including an enhanced cytokine production profile, a less-differentiated T cell phenotype with lower expression of stress and exhaustion markers, an enhanced metabolic profile and increased proliferation in TME-like conditions.

**Conclusions:** Together, these results demonstrated that co-expression of GOT2 can substantially improve the overall antitumor activity of CAR T cells by inducing broad changes in cellular function and phenotype. These data show that BOXR1030 is an attractive approach to targeting select solid tumors. To this end, BOXR1030 will be explored in the clinic to assess safety, dose-finding, and preliminary efficacy (NCT05120271).

## Introduction

Chimeric antigen receptor-based T cell (CAR T) therapeutics have revolutionized the field of oncology, and have provided substantial benefits to heavily pretreated and treatment refractory patients(1). Despite early successes targeting hematological malignancies, substantial efficacy and safety challenges limit broad application of CAR T therapy in solid tumors(2–4). One of these challenges is the immunosuppressive solid tumor microenvironment (TME) which is hypoxic, rich in myeloid-derived suppressor cells and T-regulatory cells and have increased expression of inhibitory receptors (e.g. programmed-death-ligand 1 [PD-L1])(5–9). Nutrient competition within the tumor microenvironment (TME) has been shown to limit the efficacy of checkpoint inhibitor therapies and cellular therapeutics such as CAR T therapy(10, 11).

Nutrient starvation and stress encountered in the hostile TME can impact metabolism in tumor infiltrating T cells including CAR+ T cells and indeed, these T cells have been shown to be impaired metabolically (mitochondrial mass, glycolysis) in patients(12–14). Changes in T cell metabolism including in CAR T cells have been reported to impact T cell exhaustion and persistence (15–17). Nutrient competition between tumor cells and T cells has been demonstrated to be a driving force for the development of T cell exhaustion in mouse models of cancer(18). T cell metabolism is increasingly recognized as a key contributor to multiple aspects of T cell biology, influencing effector functions(19), differentiation state and memory formation(20, 21), and survival(22). Thus, modulation of T cell metabolism could improve T cell function and potentially improve human CAR T cell therapy for cancer.

Glutamic-oxaloacetic transaminase 2 (GOT2, formerly mitochondrial aspartate aminotransferase, mAspAT) is a mitochondrial enzyme that has an essential role in glutamine metabolism as part of the malate-aspartate shuttle. GOT2 catalyzes the production of alpha-ketoglutarate (aKG) and aspartate from glutamate and oxaloacetic acid, feeding the TCA (tricarboxylic acid, Krebs) cycle and contributes to the maintenance of cellular redox balance(23, 24). Although little is known about the role of GOT2 in T cells, growing evidence suggests that GOT2 may be a key factor for optimal T cell activity. T cells from GOT2 knockout mice have impaired IFNγ production(25). In addition, both aKG and aspartate, metabolic products of the GOT2 enzymatic reaction, are involved in essential T cell processes including proliferation and differentiation(25, 26). GOT2 is expressed in activated T cells(27), and protein expression is decreased in chronically stimulated T cells(28), which leads us to hypothesize that sustained over-expression of GOT2 may improve CAR T cell function within the solid TME.

Glypican-3 (GPC3) is an oncofetal tumor antigen that is an attractive target for CAR T cell therapy due to its highly restricted expression on normal tissue and high prevalence in several adult and pediatric solid tumors(29). Glypicans are membrane-bound heparin sulfate proteoglycans, known to stimulate or inhibit growth factor activity and are expressed during development in a cell- and tissue-specific manner. GPC3 is involved in regulation of cell proliferation and apoptosis in normal development during embryogenesis and expression is largely absent in normal adult tissues. Aberrant GPC3 expression is implicated in tumorigenesis(30–32), and GPC3-positive cancers including hepatocellular carcinomas (HCC), are characterized by a highly metabolic and immunosuppressive landscape(33), and exhaustion is a common feature of tumor-resident T cells(34).

Here, we describe BOXR1030, a novel CAR T therapeutic co-expressing a GPC3- targeted CAR and exogenous GOT2. GPC3 protein expression was observed in >30% of samples from patients with HCC, squamous cell carcinoma of the lung (SCC), Merkel cell carcinoma (MCC), and liposarcoma demonstrating its attractiveness as a CAR tumor antigen target. Compared to T cells expressing CAR alone, BOXR1030 T cells demonstrated superior *in vivo* efficacy and have favorable attributes including an enhanced cytokine production profile, a less-differentiated phenotype with lower expression of exhaustion markers, and an enhanced metabolic profile. The data herein support the generation of a GPC3 targeted CAR for the treatment of select solid tumor indications and demonstrate that modulation of T cell metabolism is a promising new approach to improve CAR T cell therapy for the treatment of solid tumors.

## Materials and Methods

### Ethics

Peripheral blood mononuclear cells (PBMCs) from healthy male and female human donors were obtained from HemaCare. PBMCs were isolated from leukopaks collected in HemaCare’s FDA-registered collection centers following cGMP and cGTP collection guidelines from IRB- consented donors. All animal studies accounted for the minimal number of animals required for scientific rigor, and all studies were conducted in accordance with experimental protocols reviewed and approved by the Institutional Animal Care and Use Committee (IACUC) at Unum Therapeutics and/or SOTIO Biotech Inc.

### Chimeric Antigen Receptor Constructs and Virus Production

The GPC3 CAR construct (CAR) utilized is composed of a humanized anti-GPC3 targeted single chain variable fragment, CD8-alpha transmembrane domain, and intracellular 4-1BB costimulatory and CD3-zeta signaling domains. The BOXR1030 expression constructs encode both the GPC3 CAR and GOT2, separated by a ribosomal porcine teschovirus 2A (P2A) self-cleaving peptide sequence. Constructs were cloned into the γ-retroviral vector MP71(35). CAR and BOXR1030 encoding MP71 retroviral vector plasmids were transiently transfected into HEK293RTV cells along with packaging plasmids encoding Gag, Pol, and GaLV viral proteins. GaLV-pseudotyped retroviral supernatant was harvested, filtered, and frozen approximately 24 and 36 hours after transfection.

### T Cell Transduction and Expansion

CAR transduced T cells were generated using two methods. The first process generated research scale material, while the second process generated good manufacturing practice (GMP) analogous material. GMP analogous material was used in the safety/biodistribution study, while research scale material was used in all other evaluations.

For research scale material, commercially procured frozen PBMC from Hemacare were thawed into TexMACS media (Miltenyi Cat #130-097-196) and activated with soluble anti-CD3 antibody (OKT3; 1µg/mL), anti-CD28 antibody (15E8; 0.5µg/mL) (Miltenyi Biotec, Cat# 130- 093-387 and 130-093-375), and IL-2 (100 U/mL) (Prometheus/Novartis Cat# 0078-0495-61) for 48 hours. For retroviral transduction, T cells were concentrated to 1x10^6^ cells/mL and an equal volume of cells and retroviral supernatant were centrifuged at 1200 x g for 45 minutes. Cultures were transferred to G-Rex 10 or 100 units (Wilson Wolf Cat# 80500 and 800240M) 24 hours later and expanded in TexMACS media supplemented with IL-2 (100 U/mL) every 48-72 hours. CAR products were harvested on day 10 and cryopreserved in Bambanker cell freezing medium (Wako Pure Chemical Industries, Cat# 302-14681).

For GMP analogous material, PBMCs were isolated from peripheral blood from donors by Ficoll separation and cultured into X-Vivo 15 + 5% CTS Immune Cell SR (Lonza Cat# BE08-879H and Gibco Cat# A2596101) and activated with soluble anti-CD3 antibody (OKT3; 1µg/mL), anti-CD28 antibody (15E8; 0.5µg/mL) (Miltenyi Biotec, Cat# 130-093-387 and 130-093-375) and IL-2 (100 U/ml) (Cellgenix Cat# 1020-050) for 2 days. For spinoculation, a total of 1 x 10^8^ cells and 70 mL of retroviral supernatant were centrifuged at 2000 x g for 20 minutes. Cells were then transferred back into 250 mL conical flasks, spun for 5 minutes at 650 x g, RT and supernatant was removed. A single wash was performed, and cells were resuspended in complete culture media at a final density of 6.7×10^5^ cells/mL and transferred to 500 mL gas-permeable cell expansion bags and incubated at 37°C overnight. Cultures were transferred to gas permeable bags on day 3 and a second spinoculation was performed. Double-transduced cells were then resuspended in complete culture media at a final density of 6.7×10^5^ cells/mL and transferred to 500 mL gas-permeable cell expansion bags and incubated at 37°C + 5% CO2 as above. CAR products were harvested on day 8 and cryopreserved in 1:1 ratio of Plasma-Lyte A and CryoStor CS10.

Volume of viral vector was matched during production for both BOXR1030 and Control CARs. Specific MOIs were not used in production of BOXR1030 or the control CARs. Variability in CAR expression between BOXR1030 and Control CAR was largely donor- and batch- dependent but overall expression levels across multiple batches were comparable.

### Restimulation under hypoxic and low glucose conditions

BOXR1030 and control CAR cells were diluted with donor-matched untransduced T cells to normalize %CAR+ cells for each experiment. T cells were pre-activated with GPC3+ tumor cells in standard culture conditions (normoxia and 11 mM glucose) at an E:T ratio of 2:1 for 3 days. Co-cultured samples are mixed thoroughly by pipetting and transferred to a fresh plate with additional GPC3+ tumor cells for restimulation. Restimulation was performed for 4 days under hypoxic conditions (1.5% oxygen) or in low glucose conditions (glucose-free RPMI + 10%FBS supplemented with 2mM D-glucose). The controls for each condition were restimulated for 4 days with normal oxygen levels or in glucose-free RPMI + 10%FBS supplemented with 10 mM D-glucose as appropriate.

### Cell Lines

HepG2 (HB-8065) and Hep3B (HB-8064) hepatocellular carcinoma (HCC), PLC/PRF/5 (CRL- 8024) hepatoma and BT-20 (HTB-19) breast carcinoma cell lines were obtained from ATCC (Manassas, VA) and were maintained at 37°C and 5% CO_2_ in EMEM medium (Gibco, Cat# 670086) supplemented with 10% heat inactivated FBS. For certain studies, HepG2 cells were fixed with 4% paraformaldehyde and then stored at 4°C in RPMI 1640 (Gibco, Cat# 11875119) media supplemented with 10% heat inactivated FBS. The JHH7 (JCRB1031) HCC cell line was obtained from JCRB (Osaka, Japan) and maintained at 37°C and 5% CO_2_ in William’s E medium (ThermoFisher Scientific, Cat. # 32551) supplemented with 10% heat inactivated FBS. The HUH-7 (JCRB, JCRB0403) HCC and A375 melanoma cell lines (ATCC, CRL-1619) were maintained at 37°C and 5% CO_2_ in Dulbecco’s Modified Eagle’s Medium (DMEM) (Corning, Cat# 10-102-CV) supplemented with 10% heat inactivated FBS. SNU-398 (CRL-2233) HCC, NCI-N87 (CRL-5822) gastric adenocarcinoma, and 786-O (CRL-1932) renal cell carcinoma call lines were obtained from ATCC and maintained at 37°C and 5% CO_2_ in RPMI 1640 (Corning, Cat# 10-040-CV) supplemented with 10% heat inactivated FBS. The PC3 (ATCC, CRL-1435) prostate adenocarcinoma cell line was maintained at 37°C and 5% CO_2_ in F-12K medium (ATCC, Cat# 30-2004) supplemented with 10% heat inactivated FBS. The OV-90 (ATCC, CRL- 11732) ovarian adenocarcinoma cell line was maintained at 37°C and 5% CO_2_ in a 50% mixture of MCDB 105 (Sigma-Aldrich, Cat# M6395) and Medium 199 (ThermoFisher Scientific, 11150-059) supplemented with 15% heat inactivated FBS. The K562-ROR1 cell line was derived from K562 (ATCC, CCL-243) and maintained at 37°C and 5% CO_2_ in Iscove’s Modified Dulbecco’s Medium (IMDM) (ATCC, Cat# 30-205) supplemented with 10% heat inactivated FBS.

For retroviral production, HEK293T cells were procured from Cell Biolabs, Inc. (San Diego, CA). Cells were maintained at 37°C and 5% CO_2_ in DMEM medium containing 10% heat inactivated FBS, 2 mM L-Glutamine, and 0.1 mM nonessential amino acids. Mycoplasma testing was performed on all cell lines and confirmed to be negative.

### Immunohistochemistry

Formalin fixed paraffin embedded (FFPE) tumor tissue microarrays (TMAs) were sourced from US Biomax (Cat# FDA999w1, LV809b, LC814a, LC817b, SO302a, SO2083a, CO702d, CO489a, and OV991). FFPE blocks for HCC, liposarcoma, non-small cell lung cancer (NSCLC) and MCC were sourced by NeoGenomics Laboratories, Inc. (Aliso Viejo, CA). Sections at 5µm were stained using anti-GPC3 mouse monoclonal primary antibody (clone GC33, Ventana Medical Systems, Inc. Tucson, AZ Cat# 790-45654) on BenchMark ULTRA to detect membrane and cytoplasmic expression. After anti-GPC3 primary antibody staining, heat-induced epitope retrieval was used followed by incubation of the primary antibody for 32 minutes. Immunodetection was accomplished with OptiView DAB Detection Kit (Ventana, Cat# 760- 700). Appropriate positive and negative controls were included for each stain. All specimens had an H&E stained slide for morphological reference. Stained slides were evaluated for GPC3 expression manually by a qualified pathologist, using a standard brightfield microscope. The percent of tumor cells with GPC3 membrane and cytoplasmic staining at each intensity (0, 1+, 2+, 3+ corresponding to no staining, weak staining, moderate staining and strong staining, respectively) were recorded and an H-score (range 0-300) was calculated using the following equation:

H-score = 1*(%tumor cells with 1+ intensity) + 2*(%tumor cells with 2+ intensity) + 3*(%tumor cells with 3+ intensity)

The cutoff for positive GPC3 expression was set at H score ≥ 30 in tumor cells stained with the GPC3 IHC assay.

### Real Time PCR Analysis

CAR and BOXR1030 T cells were stimulated with fixed HepG2 target cells at a 2:1 effector-to-target cell ratio for six days. Samples were harvested and total RNA was extracted using the RNeasy Micro Kit (Qiagen, Cat# 74004), following the manufacturer’s instructions. Next, cDNA was generated using the High Capacity cDNA Archive Kit (Applied Biosystems, Cat# 4368814). For each sample from each condition, equal amounts of cDNA were used to perform quantitative PCR with custom-designed primers and probes for endogenous and exogenous GOT2. The copy number of endogenous and exogenous GOT2 was calculated by extrapolation of Ct values from a standard curve generated from a GOT2 amplicon.

### Western Blot Analysis

CAR and BOXR1030 T cells were activated with immobilized recombinant GPC3 protein (0.3 ug/mL) (R&D Systems, Cat# 2119-GP-050) over a 6-day time course. Following activation, samples were washed once with PBS and lysed in RIPA buffer containing protease inhibitor (EMD Millipore, Cat# 20-188 and 20-201). Total protein concentration was quantified using the BCA Protein Assay Kit (Fisher Scientific, Cat# 23227). Samples were normalized to a fixed amount of protein, reduced with DTT in sample loading buffer, and denatured at 95°C. Next, samples were resolved on a Bolt^TM^ 4-12% Bis Tris Plus gradient gel (ThermoFisher Scientific, Cat# NW04120BOX) and transferred onto nitrocellulose membranes. Blots were blocked in 5% non-fat milk, probed with primary antibodies at 4°C overnight, and then HRP-conjugated secondary antibodies for 1 hour at room temperature. Finally, blots were developed with Clarity- ECL HRP chemiluminescent substrate (Bio-rad, Cat# 1705060), and gel images were captured on the G:BOX Chemi XX6 gel imager (Syngene) using GeneSys software.

Western blot antibodies for GOT2 detection were rabbit anti-human GOT-2 polyclonal antibody (Origene, catalog #TA325088) and HRP-conjugated AffiniPure Goat anti-rabbit IgG (ThermoFisher, catalog #G-21234). Western blot antibodies for the GAPDH loading control were mouse anti-human GAPDH antibody (Biolegend, catalog #649202) and HRP-conjugated anti-mouse IgG (Cell Signaling Technologies, catalog #7076S).

### Flow Cytometry

Mouse anti-human antibodies to CD3 (OKT3), CD4 (OKT4; RPA-T4), CD8 (RPA-T8), CD45RA (HI100), CD27 (M-T271), IFN-gamma (4S.B3), IL-2 (MQ1-17H12), TNF-alpha (MAb11), IL-17A (BL168), CD69 (FN50), PD-1 (EH12.2H7), and TIM-3 (F38-2E2) (Biolegend) were used to evaluate T cell phenotype. Recombinant human GPC3 protein was conjugated to AlexaFluor 647 using NHS ester chemistry and used to detect CAR+ T cells. For all flow cytometry experiments, dead cells were excluded with Fixable Viability Dye eFluor 780 (Fisher Scientific, Cat# 50-169-66). For *ex vivo* phenotypic studies, rat anti-mouse antibodies to CD45 (30-F11) and CD11b (M1/70) (Biolegend) were used to exclude murine cells from analysis.

To evaluate intracellular cytokine production, CAR T cells were incubated with GPC3+ HepG2 target cells at a 1:1 effector-to-target cell ratio in the presence of brefeldin A (5 µg/mL) and monensin (2 µM) protein transport inhibitors. The co-cultures were incubated for 6 hours at 37°C and 5% CO2 prior to cell staining using the Cytofix/Cytoperm Fixation Kit (BD Biosciences, Cat# 554715/555028).

#### Enumeration of Peripheral Blood T Cells and CAR+ T Cells by Flow Cytometry

Cryopreserved peripheral blood samples from mice were thawed by incubation at 37°C for 5 minutes and washed with PBS. Red blood cells were lysed with ACK lysing buffer (Thermo Fisher, Cat# A1049201) for 5 minutes at room temperature, and then washed twice with PBS. Cells were then stained for 25 minutes on ice with an antibody cocktail of mouse anti-human antibodies to CD3 (OKT3), CD4 (OKT4; RPA-T4), and CD8 (RPA-T8) (Biolegend) and recombinant human GPC3 protein was conjugated to AlexaFluor 647 using NHS ester chemistry and used to detect CAR+ T cells. Dead cells were excluded with Fixable Viability Dye eFluor 450 (Fisher Scientific, Cat# 50-169-66).

### Aspartate Assay

T cells were thawed and rested in pre-warmed RPMI 1640 medium containing 10% FBS for 3-4 hours at 37°C and 5% CO_2_. To prepare lysates, rested T cells were washed twice with DPBS and resuspended in AST assay buffer (BioVision, Cat# K552). Lysates were homogenized with a Dounce homogenizer and subsequently deproteinated through 10 kDa Amicon filters. Intracellular aspartate was quantified using a plate-based Aspartate Colorimetric Assay Kit (BioVision, Cat# K552). Lysates (50 µL) were incubated for 30 minutes with the enzyme reaction mixture (50 µL). Absorbance was measured at an optical density of 570 nm and aspartate concentration was extrapolated from a standard curve following background subtraction.

### RNA Sequencing

CAR and BOXR1030 T cells were activated with immobilized recombinant GPC3 protein (0.3 mg/mL; R&D Systems, Cat# 2119-GP-050) for 4 and 24 hours. Samples at the 4 hour activation time point were also processed to isolate CD4+ and CD8+ T cells by positive CD4 and CD8 selection kits (Miltenyi Biotec, Cat# 130-045-101 and 130-045-201). Unstimulated T cells were processed at the 0 hour time point without any activation. Total RNA extraction and sequencing were performed by Genewiz (Cambridge, MA). RNA concentrations were calculated using the Qubit RNA Assay. Total RNA was then enriched for poly-A mRNA, followed by fragmentation. After random priming, first and second strand cDNA synthesis was performed. cDNA was then processed for end repair, 5’ phosphorylation, and dA-tailing. Adapter ligation and PCR enrichment were than completed prior to sequencing. Finally, 2x150 bp, single index RNA sequencing was performed on the Illumina HiSeq system.

The 54 FASTQ files were QC reviewed with fastQC (v) and multiQC (v). Reads were quasi-mapped using Salmon (v1.0.0) to a custom decoyed transcriptome index. The decoyed index was built using the gencode v32 transcriptome, customized with additional BOXR1030 transcripts, and decoyed using the gencode GRCh38 primary genomic assembly. Sample specific quantification files were imported into R (v3.6.1) and converted to “lengthScaledTPM” counts using the *tximport* package (v1.12.3). In order to be included in the downstream analysis, features were required to have a minimum of 20 counts in at least 6 samples (chosen due to sample numbers in key groups of interest). The feature filtered count matrix was “voom” transformed (log2(CPM)) and tested for differential expression (DE) across CAR products and longitudinally using the *limma* R (v3.40.6) package. The DE tests utilized *limma*’s *treat* analysis pipeline to test for changes in gene expression greater than 25% from the reference condition. Pathway expression scores were generated for each sample using the voom-transformed count matrix and the “ssgsea” implementation in the *GSVA* R package (v 1.32.0). Pathway scores were generated for pathways defined in *MSigDB* (v6.1) collections H, C2, C3, C5, C6, and C7. DE tests on the ssGSEA pathway expression scores utilized *limma*’s *eBayes* analysis pipeline to test for non-zero changes in pathway expression. *WGCNA* (v1.68) was used to cluster DE genes (and separately DE pathways) into groups with significantly correlated expression profiles. Gene-level modules were analyzed for pathway enrichment using *MSigDB* pathways and *fisher.test* in R for hypergeometric enrichment.

### Xenograft *in vivo* Studies

GPC3+ Hep3B and JHH7 HCC xenograft tumors were induced in 6-8 week old female NSG (NOD scid gamma, NOD.Cg-Prkdc scid IL2rg tmWj1 /SzJ, Strain 005557) mice. For the Hep3B xenograft model, 5 x 10^6^ tumor cells were inoculated subcutaneously on day 0. Animals were randomized into treatment groups at an average tumor volume of ∼100 mm^3^, and treated with 1 x 10^6^ CAR+ T cells on days 20 and 27. For the JHH7 xenograft model, 5 x 10^6^ JHH7 tumor cells were inoculated subcutaneously on day 0 and animals were treated with 5 x 10^6^ CAR+ T cells on days 8 and 15, when average tumor volume was ∼50 mm^3^. T cells were administered via intravenous injection. Tumors were measured twice weekly in three dimensions using calipers and tumor volume was calculated using the formula: 0.5 * (length * width * height). At designated time points, blood was collected from the retro-orbital sinus and cryopreserved prior to cell staining and analysis. In another JHH7 study, different doses of T cells were administered as indicated and blood was collected to monitor pharmacokinetics and toxicology parameters, including hematology and clinical chemistry to monitor organ function and potential hematologic toxicities. Animals were euthanized at Day 15 and Day 45 post CAR T cell dosing and the following tissues were collected to assess biodistribution: liver, spleen, lung, and tumor (highly vascularized tissues with expected T cell infiltration); and brain, heart, kidney and ovaries (tissues expected to have minimal T cell infiltration). Tissues were processed for histologic analysis and for detection of CAR+ T cells by qPCR.

Genomic DNA (gDNA) was extracted from snap-frozen tissues (∼10–20 mg) using DNeasy Blood and Tissue Kit (Qiagen, Cat# 69504) according to the manufacturer’s instructions, and quantified using a Quant-iT™ dsDNA Assay Kit (ThermoFisher Scientific, Cat# Q33120). DNA integrity (DIN) was analyzed on selected samples using TapeStation 4200 (Agilent), based on a scale of 1 (fully degraded) to 10 (fully intact). DNA samples (100 ng) were analyzed by qPCR and the copy number of BOXR1030 CAR scFv was calculated by extrapolation of Ct values from a standard curve generated from a synthetic CAR scFv amplicon (gBlock, IDT)

### GPC3 expression from public databases

RNA-sequencing profile of GPC3 expression in patient tumors from selected indications were generated in whole by the TCGA (The Cancer Genome Atlas) Research Network. Data were accessed using the cBioPortal for Cancer Genomics hosted by the Center for Molecular Oncology at Memorial Sloan Kettering(36, 37). Data for selected indications was downloaded from cBioPortal on January 8, 2021 and visualized using GraphPad Prism 9 software. mRNA expression is reported as RSEM (batch normalized from Illumina HiSeq_RNASeqV2) (log2(value + 1)). The RNA expression profile for GPC3 (ENSG00000147257.13) in normal human tissues was accessed from the Genome-Tissue Expression Project (GTEx) analysis release V8 (dbGaP Accession phs000424.v8.p2) on September 24, 2021 via the GTEx portal (https://gtexportal.org/home/). Data are reported as transcripts per million (TPM) and box plots are shown as the median and 25^th^ and 75^th^ percentiles. Points are displayed as outliers if values are greater than or less than 1.5 times the interquartile range.

### Statistical analyses

Data Analysis and statistical comparisons were made using Prism 8 (GraphPad Software, San Diego, CA.). Statistical comparisons were determined by the nature of data and are specifically described within figure legends. In general, comparison between two groups were determined by Student’s *t* tests, often paired for each donor. Multiple comparisons were performed by a one-way analysis of variance (ANOVA), followed by Tukey’s multiple comparison test post-hoc. Correlations were determined by Pearson’s test. Statistical methods for RNA-Seq analysis are described in the Supplementary Data. Statistical significance was defined as p<0.05.

## Results

### Expression profile of GPC3 in human normal and tumor tissue specimens

GPC3 expression has been reported at very low levels in limited normal tissues (29, 38). To confirm these results, GPC3 mRNA and protein expression was assessed in primary human non-disease tissues. GPC3 mRNA expression profile was queried from the GTEx RNAseq database representing 54 tissue sites from ∼1,000 patient samples via the GTEx portal. GPC3 mRNA was expressed at low levels predominantly in lung, adipose, tibial nerve, kidney and breast (Supplementary Figure 1A). GPC3 protein expression by IHC in a tissue microarray (TMA) (adipose and tibial nerve not represented) showed predominantly faint, cytoplasmic staining in a few tissues including heart, kidney and stomach and no expression in lung or breast (Figure 1A, Supplementary Data).

**Figure 1.**
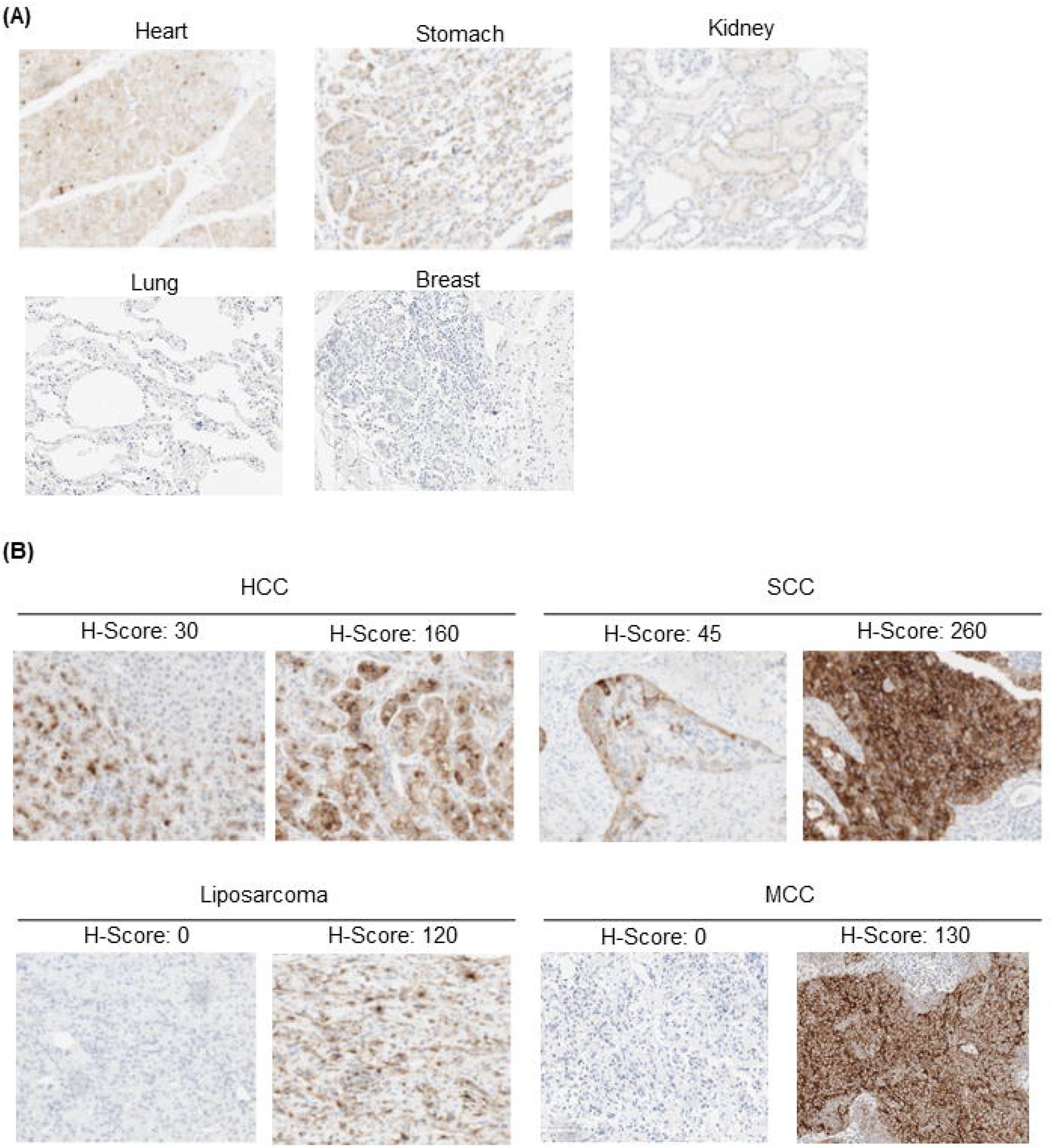
GPC3 target expression in human normal and tumor tissues by IHC. (A) Immunohistochemical staining of normal healthy TMA for GPC3 expression. Representative images (20x magnification) of different organs with different levels of GPC3 are shown (B) IHC staining of tissues from hepatocellular carcinoma (HCC), squamous cell lung cancer (SCC), Merkel cell carcinoma (MCC) and liposarcoma patients for GPC3 expression. Representative images (20x magnification) show samples with different levels of GPC3 (and H-scores).

GPC3 mRNA and protein expression was also assessed in primary human tumor samples for selected indications. RNAseq data profiling GPC3 expression in HCC, SCC, liposarcoma, colorectal adenocarcinoma (CRC), and serous ovarian cancers (OC) was accessed from the Cancer Genome Atlas Program (TCGA) and showed varying degrees of GPC mRNA expression (Supplementary Figure 1B). To determine the prevalence of GPC3 positive tumors, we performed GPC3 IHC staining of TMAs in the same indications (Supplementary Table 1A). No appreciable expression was observed in CRC or OC with <10% cases exhibiting an H-score ≥ 30. However, HCC, SCC, and myxoid/round liposarcoma (MRCLS) samples in the TMAs showed appreciable levels of GPC3 expression (H-score ≥ 30) with >20% prevalence (Supplementary Table 1A). Additionally, we performed IHC on 20-40 unique tumor samples for each of these four indications: HCC, SCC, liposarcoma, as well as, MCC(30). These additional results showed positive GPC3 staining (H-score ≥ 30) in >30% cases (Figure 1B, Supplementary Table 1B). Overall, HCC, SCC, MRCLS, and MCC had a prevalence of 69%, 33%, 33%, and 70% respectively. These data support the generation of an GPC3 targeted CAR for the treatment of these solid tumor indications.

### BOXR1030 T cells co-express exogenous GOT2 and a GPC3-targeted CAR

We designed a 2^nd^ generation CAR containing a humanized GPC3-targeting scFv (derived from codrituzumab, also known as GC33), 4-1BB costimulatory domain, and a CD3ζ signaling domain. Codon optimized GOT2 was co-expressed in the same construct using a P2A sequence to evaluate the impact of modulation of T cell metabolism on CAR T cell function, referred to as BOXR1030 herein (Figure 2A).

**Figure 2.**
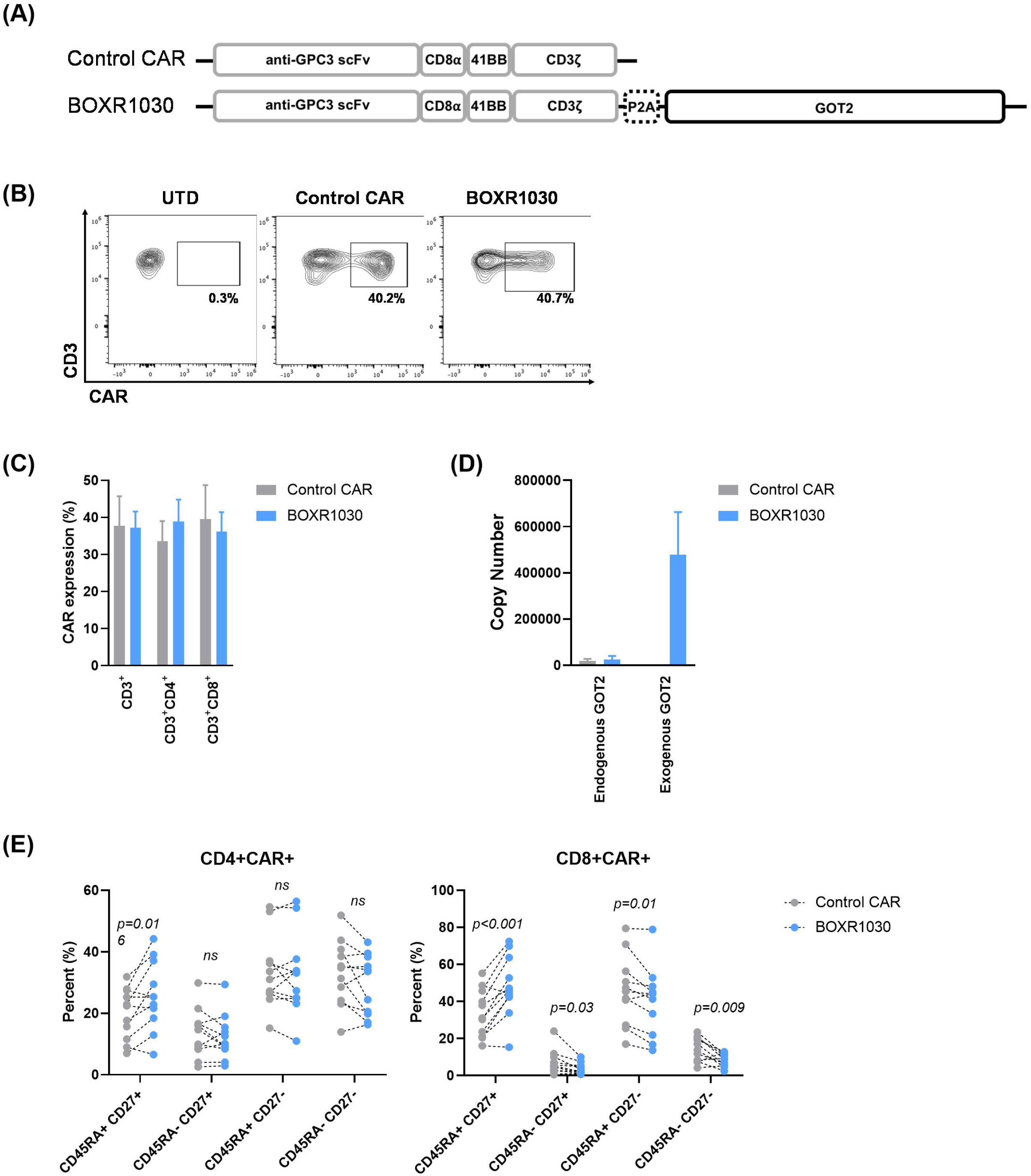
Co-expression of GPC3-targeted CAR with exogenous GOT2 in BOXR1030 T cells. (A) Depiction of expression construct and domains for the GPC3 CAR and BOXR1030 used in studies. (B) Representative surface expression of the anti-GPC3 scFv following transduction, measured on day 9 by flow cytometry from a single donor. (C) Summarized CAR expression by flow cytometry for n=5 healthy donors. (D) mRNA copy number of endogenous and exogenous (codon optimized) GOT2 measured by qPCR (n= 3 donors). (E) The frequency of CD45RA+ CD27+, CD45RA+ CD27-, CD45RA- CD27+, and CD45RA- CD27- T cells in the indicated subsets at baseline (n=11 healthy donors; no stimulation). P-values were determined by paired t-test. Data represented as mean+/- standard deviation (SD).

BOXR1030 T cells or control T cells expressing the GPC3-targeting CAR alone, were generated by activating normal human donor PBMCs with anti-CD3 and anti-CD28 and transducing with gamma retrovirus. T cells were assessed after 10 days for CAR and GOT2 transgene expression. GPC3 CAR was expressed at similar levels in both control CAR and BOXR1030 CD3+ T cells (Figure 2B and C). CAR expression was also equivalent in CD4+ and CD8+ T cell populations in both control and BOXR1030 T cells (Figure 2C). To distinguish between endogenous and exogenous GOT2, qRT-PCR primers were designed to specifically detect the wild-type and codon optimized GOT2 sequences. Endogenous GOT2 mRNA expression was detected at similar levels in both control CAR T cells and BOXR1030 cells, whereas exogenous GOT2 mRNA levels were only detected in BOXR1030 T cells and at levels significantly higher than endogenous GOT2 mRNA (Figure 2D).

GOT2 protein levels were measured by Western blot. Because endogenous and exogenous protein have the same amino acid sequence, they are indistinguishable. Despite detecting high levels of mRNA expression in BOXR1030 T cells, there was no apparent difference in protein expression at baseline compared to control CAR T cells (Supplementary Figure 2A). However, when T cells were stimulated with plate bound GPC3 antigen for six days, an increase in GOT2 protein expression was observed in a time dependent manner in BOXR1030 T cells, while GOT2 expression in control CAR T cells remained unchanged (Supplementary Figure 2A). Similarly, exogenous GOT2 mRNA expression increased in BOXR1030 T cells following stimulation and directly correlated with an increase in GPC3 CAR scFv mRNA expression (Supplementary Figure 2B-D). Taken together, these data demonstrate GPC3 CAR and GOT2 expression in BOXR1030 T cells that is maintained throughout 7 days post T cell activation.

We next evaluated T cell phenotype in CD8+CAR+ and CD4+CAR+ T cells from BOXR1030 and control CAR T cells. CD45RA and CD27 are surface markers associated with T cell memory and differentiation state. BOXR1030 T cells had a greater frequency of CD8+CAR+ T cells with a less-differentiated naïve phenotype (CD45RA+CD27+) relative to control CAR T cells and corresponding decreases in more differentiated central memory (CD45RA-CD27+), effector memory (CD45RA-CD27-), and effector memory-like (CD45RA+CD27-) populations (Figure 2E). A similar, but less pronounced trend was observed in CD4+CAR+ T cells with a significant increase in the less-differentiated CD45RA+CD27+ population (Figure 2E). Taken together these results suggest that BOXR1030 T cells may have a less differentiated and therefore more favorable phenotype compared to T cells expressing CAR alone.

### Exogenous GOT2 overexpression enhances intracellular aspartate and aspartate aminotransferase (AST) activity

GOT2 catalyzes the production of aspartate and aKG (23, 24) (Figure 3A). To confirm function of exogenously expressed GOT2, we measured aspartate and aKG levels in control CAR and BOXR1030 T cells. aKG levels were below the limits of detection of the assay (data not shown), however we were able to detect a significant increase in intracellular aspartate in BOXR1030 T cells relative to control CAR T cells (Figure 3B). AST enzymes include GOT1 and GOT2 and catalyze the reversible conversion of oxaloacetate (OAA) and glutamate into aspartate and αKG, respectively. To further confirm activity of GOT2, BOXR1030 and CAR control T cells were activated with GPC3+ Hep3B target cells for 8 days to evaluate the contribution of GOT2 to AST activity. Although magnitude of activity varied between experiments, overall greater AST activity was observed in BOXR1030 T cells relative to controls (Figure 3C). In summation, exogenous expression of GOT2 in BOXR1030 T cells results in increased AST activity and intracellular aspartate consistent with an enhanced metabolic phenotype compared to T cells expressing CAR alone.

**Figure 3.**
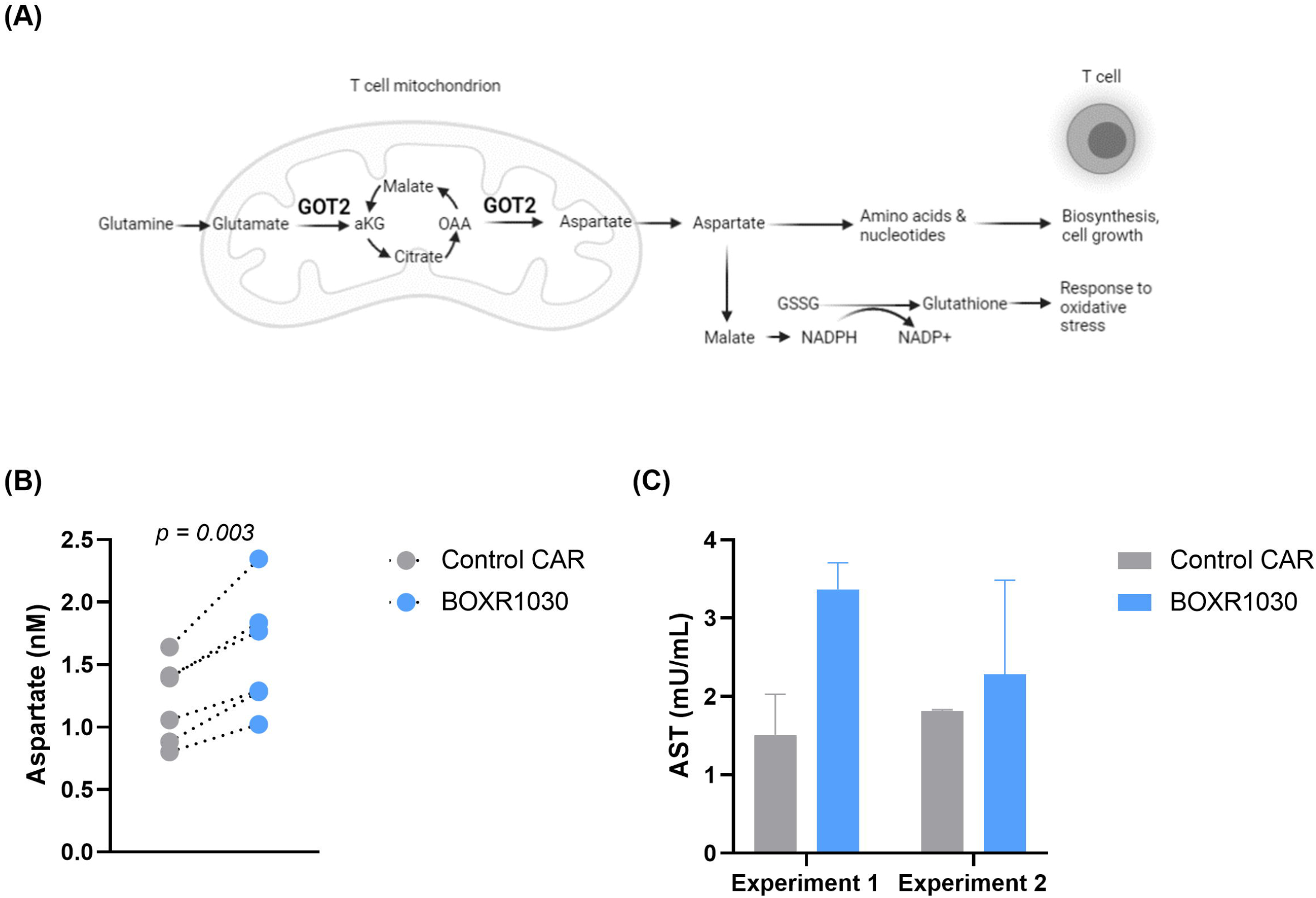
AST enzymatic activity in BOXR1030 CAR T cells. (A) Schematic describing the role of GOT2 in mitochondrial conversion of glutamate to aKG and OAA to aspartate. Image created with biorender.com. (B) Intracellular aspartate levels were measured using a plate-based colorimetric assay comparing BOXR1030 and control CAR T cells under non-stimulated conditions. (C) BOXR1030 and control CAR T cells were activated for 8 hours with GPC3+ Hep3B target cells and evaluated for AST activity. Data represented as mean +/- SD of two technical replicates.

### BOXR1030 has enhanced in vitro activity under TME-like stress conditions

*In vitro* activity of BOXR1030 and CAR control T cells was evaluated in co-culture with tumor cell lines. Cytokine secretion, cytotoxicity, and T cell proliferation were measured in response to a panel of cell lines expressing high, medium, and low levels of the GPC3 target antigen to determine whether *in vitro* activity was dependent on antigen expression level (Figure 4A and B). BOXR1030 T cells performed similarly to control CAR T cells in all assays, indicating comparable CAR function in the presence of GOT2 transgene under standard *in vitro* assay conditions (Figure 4C-H). IFN-gamma release was observed against all target expressing cell lines regardless of high, medium, and low GPC3 expression levels (Figure 4C). IL-2, TNF-alpha, and IL-17A release was observed with high and medium GPC3 expressing lines, and decreased levels were observed with the low GPC3 expressing target cell lines (OV90 and SNU398) (Figure 4D-F). PLC/PRF/5 cell line has overall low GPC3 expression, however expression is heterogenous in this cell line with populations ranging from negative to high expression (Figure 4B), which may explain higher levels of activity of response with this cell line.

**Figure 4.**
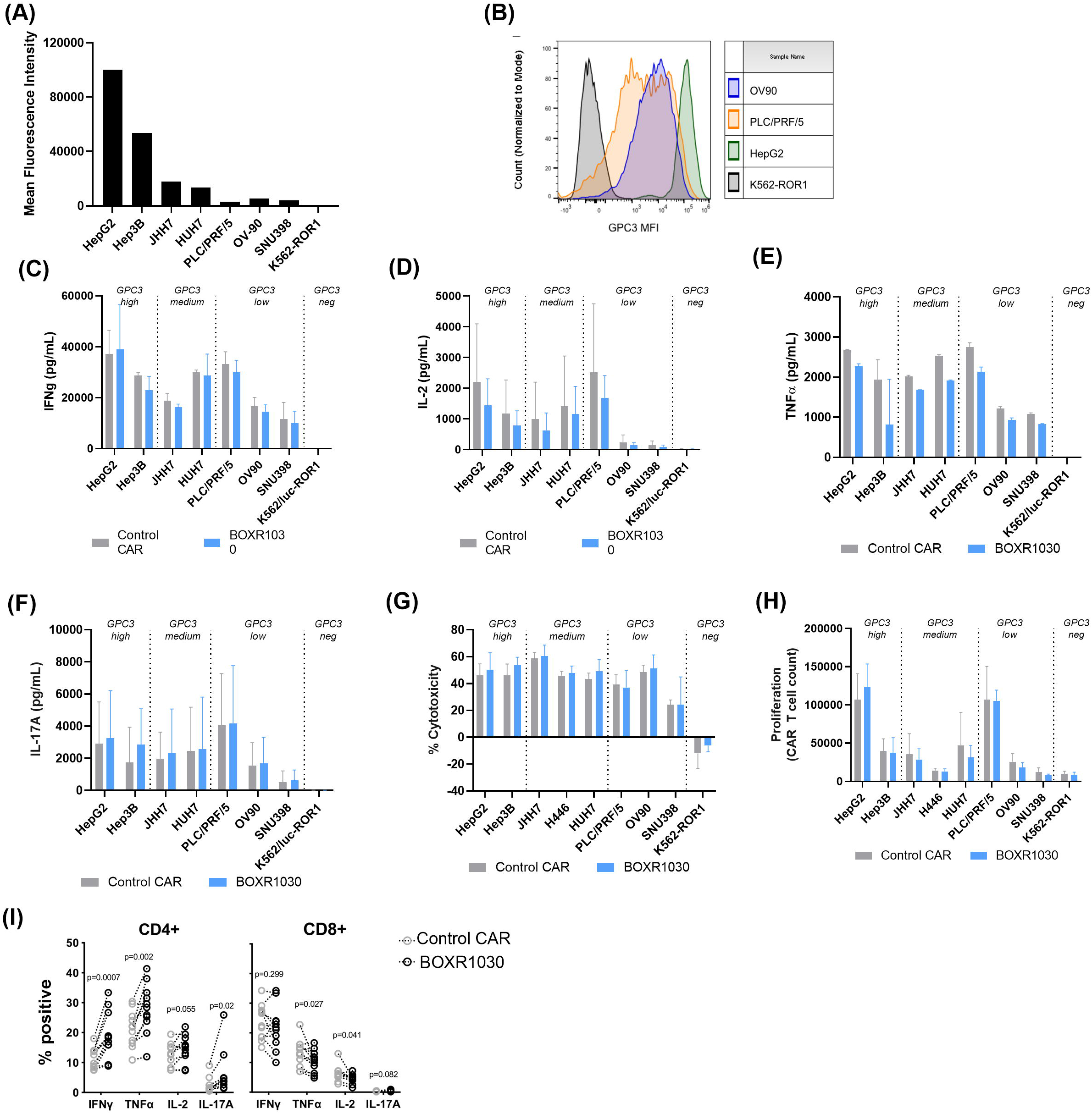
*In vitro* BOXR1030 activity. (A) Expression of GPC3 target antigen on target cell lines were measured by flow cytometry and represented as median fluorescence intensity (MFI). (B) Histogram plots of GPC3 MFI for selected cell lines. (C-F) T cell cytokine release of BOXR1030 and control CAR T cells was measured by ELISA following co-culture with target cell lines for 24 hours. Results for IFN-g (C) and IL-2 (D) are the averages across 3 donors, and error bars indicate SD. Results for TNF-a (E) and IL-17A (F) are for a single donor, and error bars indicate SD between technical replicates. (G) Target cell cytotoxicity was measured with a luciferase-based assay, and percent cytotoxicity was normalized to samples treated with untransduced T cells following 24 hours of co-culture (n=3 donors, error bars indicate SD). (H) T cell proliferation was evaluated after a 7 day incubation with target cell lines. CAR+ T cell counts were measured by flow cytometry. (n= 3 donors, error bars indicate SD). (I) Intracellular cytokine levels were measured by flow cytometry in CD4+ and CD8+ T cells (n=10, error bars indicate SEM).

Based on the slightly different cell phenotypes observed in CD4+ and CD8+ BOXR1030 cells (Figure 2E), we also evaluated the ability of CD4+ and CD8+ CAR+ T cells to produce TNF-α, IFN-γ, IL-2, and IL-17A following stimulation with GPC3+ target cells by intracellular cytokine staining. Relative to control CAR CD4+ T cells, we observed a significant increase in the proportion of BOXR1030 CD4+ T cells that produced IL-17A, IFN-γ, or TNF-α (Fig 4I). These changes were more pronounced within the CD4+ subset, and within the CD8+ subset, we observed no difference for either IFN-γ or IL-17A and a slight decrease in TNF-α+ and IL-2+ CD8+ BOXR1030 T cells (Fig 4I). This suggests increased numbers of pro-inflammatory Th1 and Th17 CD4+ T cells which can, in addition to augmenting anti-tumor cytotoxic responses by CD8+ T cells, mediate direct anti-tumor activity by the production of cytokines (34, 35).

T cell cytolytic activity was comparable across all target antigen densities (Figure 4G), whereas T cell proliferation was greater in high and medium GPC3 expressing lines compared to low expressing lines (Figure 4H). A threshold of GPC3 expression for activation of BOXR1030 T cells was not identified, however diminishing levels of activity were observed in low GPC3 antigen cell lines. Importantly, no reactivity was observed against a panel of GPC3 negative cell lines confirming specificity of the GPC3 CAR for its target antigen (Supplementary Figure 3A- E).

To further test BOXR1030 T cells *in vitro*, CAR T cells were assessed under conditions of chronic antigen stimulation in the presence of limited oxygen or nutrients to mimic the solid TME. BOXR1030 or control CAR T cells were co-cultured with either Hep3B or JHH7 target cells under TME-stress conditions and proliferation was measured by geometric mean fluorescence intensity of the intravital fluorescent dye, CellTrace Violet (CTV), where signal decreases with increased cell division. BOXR1030 T cells had consistently improved proliferation when restimulated with either Hep3G or JHH7 target cells under hypoxic conditions compared to control CAR T cells (Figure 5A). Similarly, BOXR1030 T cells also outperformed control CAR T cells in response to chronic stimulation in low glucose conditions (Figure 5B). Taken together, these data suggest that BOXR1030 T cells may have a proliferative advantage within the solid TME.

**Figure 5.**
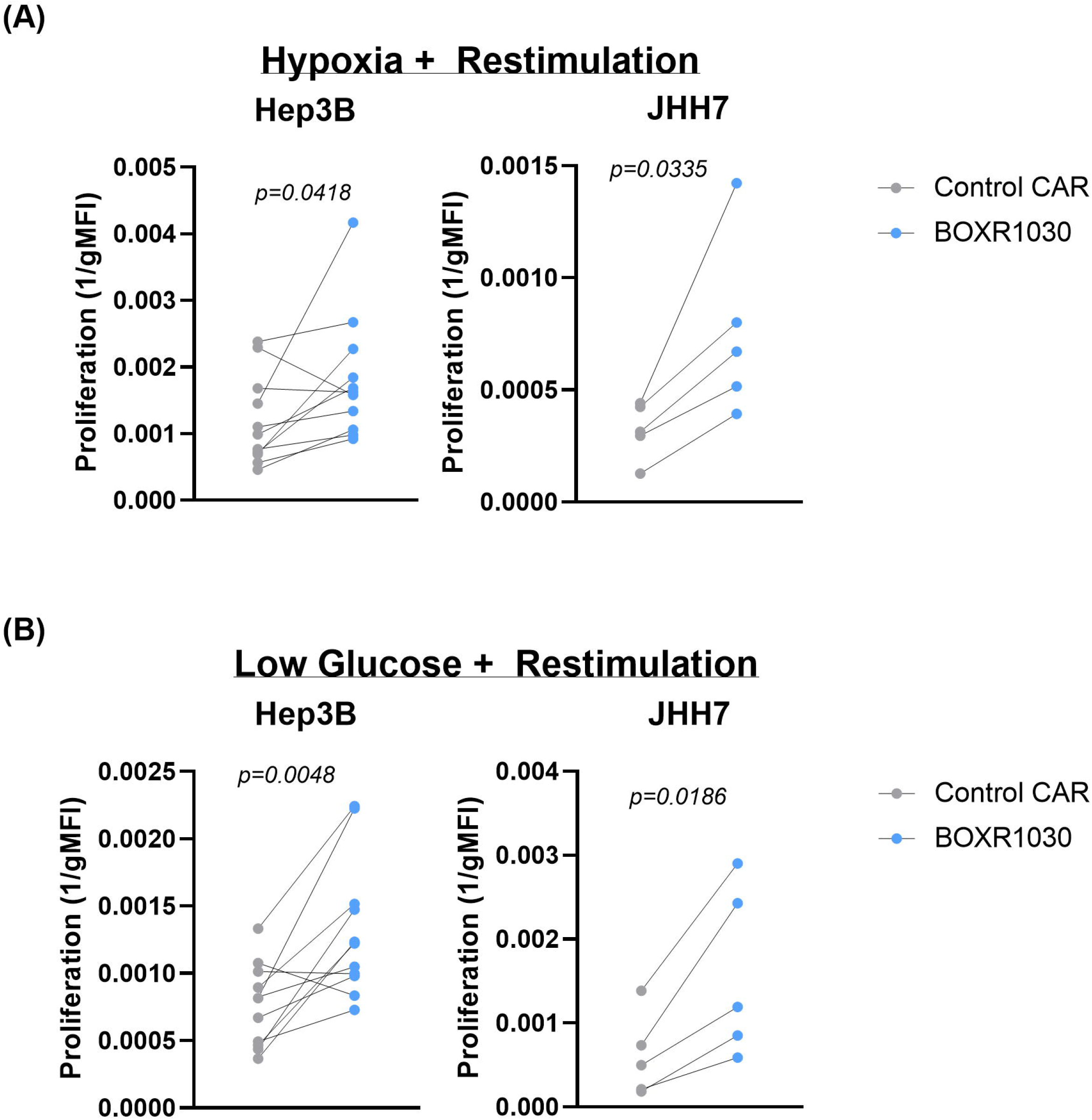
*In vitro* activity of BOXR1030 under conditions simulating the solid tumor microenvironment. BOXR1030 or control CAR T cells were repeat stimulated with GPC3+ target cell lines on day 0 and on day 3 in hypoxic conditions (A) or low glucose conditions (B). (A and B) The gMFI of CellTrace Violet (CTV) was measured for BOXR1030 and control CAR stimulated with target cells on day 7 in the indicated culture conditions and proliferation was plotted as 1/gMFI. (n=5 for JHH7 stimulated conditions and n=11 for Hep3B stimulated conditions, statistical analysis was performed using a 2-tailed paired t-test, and p-values <0.05 were considered statistically significant).

### Exogenous GOT2 overexpression alters the transcriptional profile of CAR T cells

We next evaluated the transcriptional profile of BOXR1030 T cells compared to CAR T cells at baseline and 4 and 24 hours after stimulation with immobilized recombinant GPC3 protein. T cell activation makers *CD69*, *CD154 (CD40LG)*, *IL2RA (CD25)*, and *PDCD1 (PD1)* mRNA were all elevated at 4 hours compared to baseline and returned to levels at or below baseline within 24 hours except for *CD69* which remained elevated 24 hours post stimulation. There were no differences in expression of the selected activation markers between control CAR T cells and BOXR1030 T cells (Supplementary Figure 4A).

Transcriptional differences were observed across time points following stimulation (Supplementary Table 3). Relative to the control CAR T cells, we observed significantly lower levels of genes involved in the response to cellular stress, notably *HSP90AA1, HSPE1, HSPA1A, HSPB1* and the related genes *AHSA1* and *FKBP4* (Supplementary Figure 4B-E; and Supplementary Table 3) at baseline and following activation. We also observed lower levels of the transcription factor *TOX2* in BOXR1030 T cells relative to CAR T cells. *TOX2* is a member of the TOX family of transcription factors which have recently been implicated in the development of T cell exhaustion, and knockdown of TOX2 in T cells improved their antitumor activity and survival(39). Interestingly, we also observed down regulation of *SOX4*, which has been shown to be down regulated in TOX knockout T cells(40). These data suggest that BOXR1030 cells exhibit a transcription profile consistent with reduced cellular stress and exhaustion compared to control CAR T cells.

We also observed increases in RNA levels of CD4+ T cell subset Th17 related genes such as *IL-17A/F*, as well as, CD8+ effector memory related genes such as chemokine receptor *CX3CR1*(41) in BOXR1030 cells. We extended our studies to evaluate BOXR1030 CD4+ and CD8+ T cell subsets separately. RNA levels of genes in the heat shock protein family and exhaustion related genes *TOX2* and *SOX4* were lower in both BOXR030 CD4+ and CD8+ T cell subsets (Supplementary Figure 4F and G; and Supplementary Table 2); however, select changes were restricted to a given T cell subset. For example, increases in *IL-17A/F* were restricted to the CD4+ subset, along with increased *RORC* which is a Th17 master transcription factor, implicating a possible enrichment of Th17 CD4+ T cell subsets in BOXR1030 T cells which is consistent with the previous intracellular cytokine staining data (Figure 4I). The significance of Th17 cells in anti-tumor responses has been complicated by conflicting reports. However, in the context of adoptive cell therapy such as CAR T cells, Th17 cells have exhibited persistence *in vivo*, displayed resistance to activation-induced cell death, and possessed robust recall responses which could support a more potent anti-tumor response(42).

We next examined changes in pathways between CAR and BOXR1030 at baseline and after 4 hours of activation, using single sample gene set enrichment analysis (ssGSEA). Consistent with our differential gene expression analysis, we observed downregulation of pathways associated with responses to metabolic, environmental, and oxidative stress. Notably, we observed a reduction in the pathway enrichment for genes associated with protein refolding and mTOR signaling (Supplementary Figure 4H and I). mTOR serves as a central regulator of cell metabolism, survival and response to stress, and plays essential roles in T cell memory formation(43). Previous reports have shown that reduction in mTORC1 activity leads to decreased glycolysis and prevents T-cell differentiation which can result in less-differentiated stem cell memory T cell phenotypes(44). Taken together, gene expression changes imply that BOXR1030 T cells may have reduced levels of cellular stress and T cell exhaustion compared to CAR alone control T cells.

### GOT2-expressing CAR T cells have substantially improved in vivo antitumor activity associated with reduced inhibitory receptor expression

We next evaluated the *in vivo* efficacy of CAR T cells. In order to meaningfully compare control CAR and BOXR1030 T cells, we developed two stringent xenograft tumor model systems. Our first model system, Hep3B HCC xenograft model, evaluates the intrinsic ability of small numbers of CAR T cells to expand, function, and persist through chronic stimulation and has been termed an “*in vivo* stress test”(45). When low doses of 1 x 10^6^ CAR-positive T cells were administered twice, the CAR has little to no antitumor activity (Figure 6A). In contrast, 1 x 10^6^ BOXR1030 given twice, led to sustained, durable regressions (∼100 days) for all mice treated and was well-tolerated (Figure 6A). This improved antitumor activity was associated with greater T cell expansion and long-term persistence of BOXR1030 T cells in the blood (Figure 6C), an observation consistent with a less differentiated T cell phenotype and reduced T cell exhaustion. We also evaluated a JHH7 xenograft model with intrinsic resistance to CAR function. With 5 x 10^6^ CAR+ cells administered weekly for 2 weeks, BOXR1030 demonstrated dose-dependent peripheral blood expansion and persistence (not shown) which correlated with antitumor activity against GPC3+ xenografts (Figure 6B).

**Figure 6.**
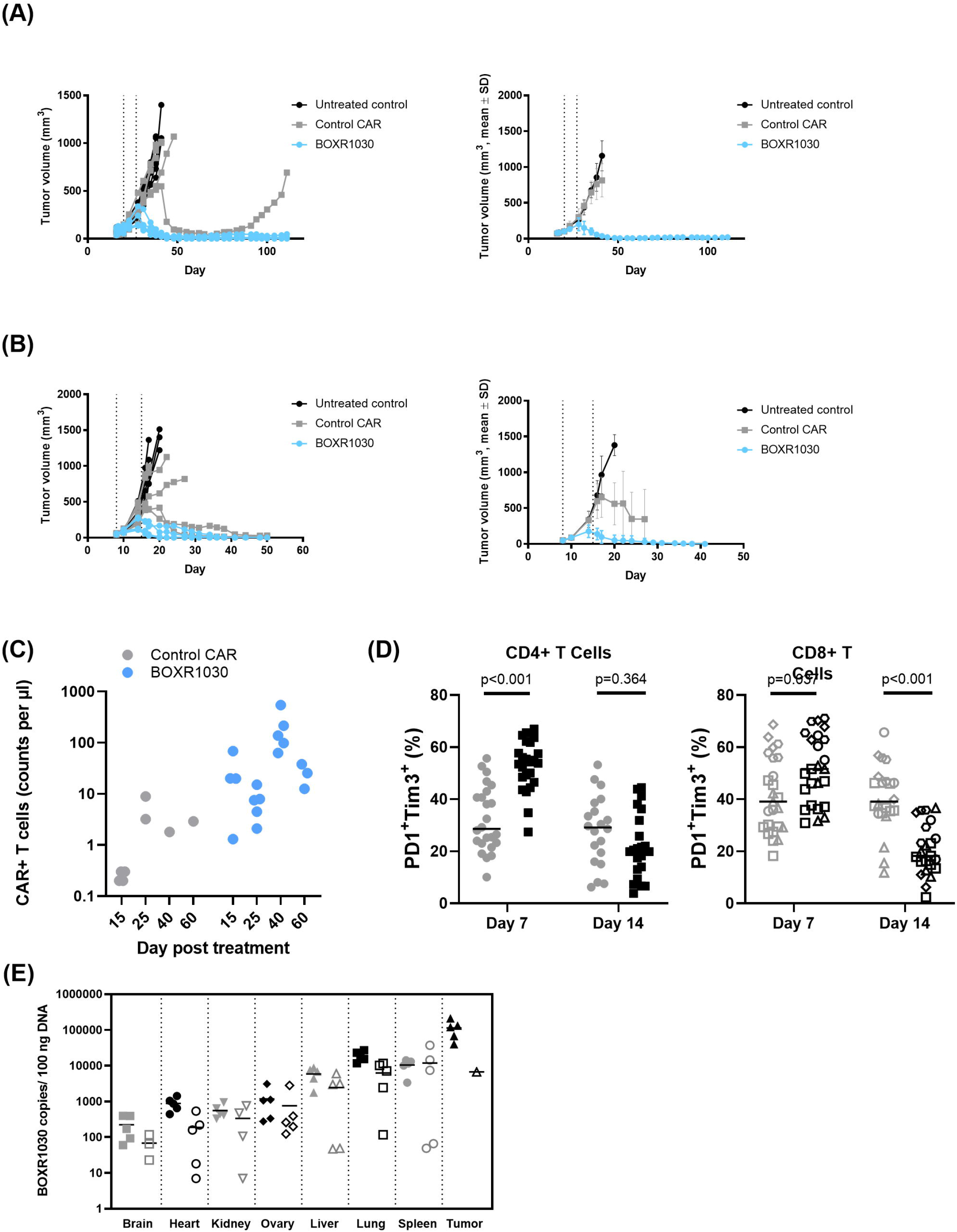
*In vivo* anti-tumor activity of BOXR1030 T cells. (A) Plot on the right shows Hep3B tumor-bearing model (mean tumor volume 108.7±34.1mm3) NSG mice were treated with two weekly doses of 1 x 10^6^ CAR+ control or BOXR1030 T cells each (total dose of 2 x 10^6^ CAR+ cells) (dosing days indicated by arrows) and tumor volumes were measured over the course of 110 days. Plot on the left shows data for individual mice and plot on the right shows mean data. Mean tumor volume plots are discontinued when less than 50% of group is remaining. (B) Plot on the right shows JHH7 tumor-bearing mice (mean tumor volume 49.8±7.2mm3) were treated with two weekly doses of 5 x 10^6^ CAR+ control or BOXR1030 T cells each (total dose of 10 x 10^6^ CAR+ cells) (dosing days indicated by arrows) and tumor volumes were measured out to 50 days. Plot on the left shows data for individual mice and plot on the right shows mean data. Mean tumor volume plots are discontinued when less than 50% of group is remaining. (C) CAR+ T cells were measured in peripheral blood of Hep3G tumor bearing mice on days 15, 25, 40 and 60 post T cell treatment and data are reported as counts per ul of blood. (D) Percent PD1+TIM3+CD4+ and PD1+TIM3+CD8+ tumor infiltrating T cells were measured by FACS on days 7 and 14 following T cell administration. (E) Biodistribution of BOXR1030 T cells was measured in mouse tissues by qPCR at days 15 and 45 post treatment. Data are reported as BOXR1030 copies / 100ng DNA.

To better characterize the T cell response leading to improved activity of BOXR1030 T cells, we selected the JHH7 xenograft model based on pilot adoptive transfer studies where we observed a relatively large influx of T cells into the tumor in the CAR group, allowing sufficient recovery for flow cytometry analysis. Prolonged exposure to the solid tumor microenvironment leads to metabolic and functional T cell exhaustion, an inhibitory program that limits the efficacy of T cells in the TME(46). This functional exhaustion is often phenotypically marked by the upregulation and subsequent co-expression of multiple inhibitory receptors. To that end, we characterized PD-1 and TIM-3 (T cell immunoglobulin and mucin domain-containing protein 3) inhibitory receptor expression on tumor infiltrating T cells at an early (day 6 or 7) time point where upregulation of PD-1 and TIM-3 are expected with T cell activation, and at a late (day 13 or 14) time point where co-expression of these receptors negatively regulate T cell activation and correlate with T cell exhaustion. BOXR1030 T cells demonstrated a higher frequency of CD4+ and CD8+ T cells co-expressing PD-1 and TIM-3 at the early time point, consistent with an activation phenotype (Figure 6D). At the later time point, CD8+ T cells of the BOXR1030 group, and not CD4+ T cells, demonstrated a significant decrease in co-expression of PD-1 and TIM-3 (Figure 6D). Importantly, expression of PD-1 and TIM-3 on CD8+ T cells, as well as on CD4+ T cells, were positively correlated to tumor volume on day 14 and supports the concept that sustained inhibitory receptor expression is associated with lack of antitumor activity. Taken together, limited expression of inhibitory receptors and sustained antitumor efficacy suggest that exogenous GOT2 overexpression may allow GPC3-targeted CAR T cells to resist T cell exhaustion when exposed to the solid tumor microenvironment.

### In vivo safety, pharmacokinetics and biodistribution of BOXR1030

A non-GLP (Good Laboratory Practice) safety and biodistribution study of GMP- analogous BOXR1030 was conducted at Charles River Research Laboratories (Durham, N.C.). Mice bearing established subcutaneous JHH7 GPC3+ xenografts were treated with a dose of 5 x 10^6^ BOXR1030+ T cells on days 1 and 8. There was no indication of toxicity; hematology and clinical chemistry values were in normal range, and no tissue toxicity was detected by histopathologic analysis (data not shown). Peripheral blood pharmacokinetics analyzed by flow cytometry show a peak of circulating BOXR1030+ T cells at day 15 (∼50 cells/uL blood) with about a 5-fold decrease in circulating BOXR1030+ T cells (∼10 cells/uL) by day 45 (data not shown). Next, we evaluated tissue distribution of BOXR1030+ T cells was analyzed by qPCR, expressed as copies per 100 ng extracted DNA. Minimal BOXR1030+ T cells were detected in brain, heart, kidney and ovaries with reduced counts at the later time point, with an average of 708 copies/100 ng DNA at day 15 and 368 copies/100 ng DNA on day 45. The more highly vascularized tissue group (liver, lung and spleen) had a greater distribution of BOXR1030+ T cells, which also demonstrated clearance over the time points with an average of 11,716 copies/100 ng DNA at day 15 and 6,840 copies/100 ng DNA on day 45. BOXR1030+ T cells were target specific, with approximately 10-fold higher accumulation in tumor tissue compared to the liver, spleen, and lung on day 15 (average 112,963 copies/100 ng tumor DNA). Only one animal had tumor tissue on day 45, and copy number in the tumor tissue at that time was comparable to liver, lung and spleen. Figure 6E shows BOXR1030+ T cells tissue and tumor distribution at day 15 and 45 post treatment.

The diminishing counts of BOXR1030 in tissue and blood over the time course of the study demonstrate that BOXR1030 does not continue to expand in the absence of GPC3+ target, supporting the target specificity and absence of antigen-uncoupled proliferation of BOXR1030. Thus, there was no indication of toxicity due to BOXR1030, and there was no indication of off-target expansion in normal mouse tissues. BOXR1030 expansion in GPC3+ tumor tissues was observed, demonstrating the target-specificity of BOXR1030.

## Discussion

In this study, we characterize BOXR1030 cells expressing GPC3-targeted CAR and GOT2 for the treatment of solid tumor indications and demonstrate that modulation of T cell metabolism can improve CAR T function in an immunosuppressive TME. GPC3 target expression analysis shows protein was expressed in >30% of cases in HCC, SCC, MRCLS and MCC solid tumor indications, and where applicable is consistent with mRNA expression in these indications. Normal tissue mRNA expression is consistently low across nearly all tissues assessed, with highest levels noted in lung, adipose, and tibial nerve; however, GPC3 protein expression by IHC confirms low and predominantly cytoplasmic localization in normal tissues making GPC3 an attractive CAR tumor antigen target.

Solid tumors are characterized by a highly metabolic and immunosuppressive microenvironment(33), and T cell exhaustion is a common feature of tumor-infiltrating T cells(34). Loss of oxidative metabolism due to reduced mitochondrial biogenesis in T cells within the TME has been observed (46). In addition, direct ligation of PD-1 on T cells can lead to broad metabolic reprogramming associated with impairment in glycolysis and amino acid metabolism(47). Activated T cells require large amounts of glucose(48) and amino acids to support biosynthesis and proliferation (19,22,25,49). Competition for nutrients within the TME has been described as a primary factor resulting in loss of T cell function(18). Thus, metabolic impairment is emerging as a key mechanism of T cell immunosuppression exhaustion within the TME.

GOT2 is a well-described mitochondrial enzyme that supports several aspects of cellular metabolism. It is expressed in activated T cells(27), and is downregulated in chronically stimulated T cells(28). In this study, we tested the effect of overexpression of exogenous GOT2 in combination with GPC3 CAR (BOXR1030 T cells) on *in vitro* and *in vivo* CAR T cell phenotype and function compared to GPC3 CAR control T cells. BOXR1030 T cells had increased levels of aspartate and AST activity confirming GOT2 functionality in these cells. In comparing BOXR1030 cells to CAR alone control cells, overall cytokine production, proliferation, and cytotoxicity under standard conditions were similar *in vitro*. However transcriptomic and intracellular protein analysis showed increased levels of Th1/Th17 related cytokines (IFNγ+, IL-17+) in CD4+ T cells expressing BOXR1030 compared to control CAR cells suggesting enhanced cytokine production profile *in vitro*. The differences in cytokine results by intracellular protein staining and ELISA bulk protein analysis may be due to masking of subset-specific differences in overall T cell secreted protein levels or the different timepoints assessed. Increased IFNγ and IL-17 expression suggests enrichment of Th1/Th17 in BOXR1030 cells which could support a more potent anti-tumor response(42). The GOT2 byproduct aKG has been reported to preferentially expand either Th1(21) or Th17(50) T cells through epigenetic modulation of gene expression. In addition, BOXR1030 cells had a less differentiated CD27+ CD45RA+ phenotype which can improve *in vivo* persistence, while also demonstrating superior anti-GPC3 function *in vitro* in TME-like conditions (hypoxia, low glucose, and chronic stimulation). Transcriptomic analysis of BOXR1030 cells also illustrated reduced expression of pathways associated with cellular stress such as protein refolding and mTOR signaling compared to CAR control only cells. Reduced cell stress and exhaustion, enhanced production of cytokines and a less-differentiated phenotype have been shown to be associated with activity of CAR T therapies and are likely key mechanistic changes responsible for the improved antitumor activity of BOXR1030 T cells relative to CAR T cells.

*In vivo,* we observed improved anti-tumor activity in two xenograft models associated with enhanced BOXR1030 expansion and persistence compared to control CAR T cells. Significantly reduced levels of PD-1 and TIM-3 expression were also observed on tumor-infiltrating BOXR1030 T cells—a key readout associated with tumor volume in our models. It should be noted that T cell dose-dependent body weight loss consistent with graft-versus host disease (GVHD) was observed at late time points in animals experiencing durable complete regressions (BOXR1030) and is likely attributed to xenogeneic reactivity of human T cells in the NSG recipient. Xeno-GVHD has been found to be dependent upon T cell dose and PBMC donor across multiple T cell therapeutics evaluated in other experiments. We did not observe any indication of toxicity or off-target expansion of BOXR1030 in normal mouse tissues, while target-specificity of BOXR1030 was observed with expansion in the GPC3+ tumor tissues.

Here, we demonstrated a mechanistic role for exogenous expression of GOT2 in the improvement of CAR T cell metabolism that was associated with changes in T cell phenotype, cytokine profile, expression of inhibitory markers, and substantially improved antitumor efficacy of a GPC3 CAR in aggressive solid tumor xenograft models. Together, these data show that BOXR1030 is an attractive approach to targeting select solid tumors. To this end, BOXR1030 will be explored in the clinic to assess safety, dose-finding, and preliminary efficacy (NCT05120271).

## Acknowledgements

The authors would like to recognize Andrew Lysaght at Immuneering for assistance with RNA-Seq analysis. The authors would like to recognize Jessica Wan for the development of the exogenous GOT2 PCR assay, and Ruchi Newman for consultation on the RNA-Seq data. The authors would like to thank all of their colleagues at Unum Therapeutics and SOTIO Biotech Inc. for feedback and critical review of data throughout the project. In particular, the authors would like to thank Jessica Sachs, Casey Judge, Amy Holbrook, and Matt Osbourne for critical review of earlier versions of the manuscript.

