## Supplementary Data for "BOXR1030, an anti-GPC3 CAR with exogenous GOT2 expression, shows enhanced T cell metabolism and improved antitumor activity"

**Results**

GPC3 IHC expression in non-diseased adult tissue

Thirty-three non-diseased adult tissue types from 3 different donors were stained and scored. Faint cytoplasmic staining for GPC3 was noted in 1 adrenal gland specimen, in 2 pancreas cores and 1 heart specimen. Staining was also reported in epithelial cells of 2 stomach cores and the ganglion cells in the 3 colon cores. Tubular epithelium staining was reported in 2 kidney cores. These results showed predominantly faint, cytoplasmic staining and are consistent with literature reported GPC3 expression profiles(1,2). There have also not been any reported off-tumor on-target toxicities in another GPC3 CAR clinical trial(3).

1.	Baumhoer D, Tornillo L, Stadlmann S, Roncalli M, Diamantis EK, Terracciano LM. Glypican 3 expression in human nonneoplastic, preneoplastic, and neoplastic tissues: a tissue microarray analysis of 4,387 tissue samples. Am J Clin Pathol [Internet]. Am J Clin Pathol; 2008 [cited 2021 Oct 8];129:899–906. Available from: http://www.ncbi.nlm.nih.gov/pubmed/18480006

2.	Al-saraireh Y, Alrawashdeh F, Al-shuneigat J, Alsbou M, Alnawaiseh N, Al-shagahin H, et al. Screening of Glypican- 3 Expression in Human Normal versus Benign and Malignant Tissues: A Comparative Study Glypican- 3 expression in cancers. Biosci Biotechnol Res Asia [Internet]. 2016 [cited 2021 Oct 8];13:687–92. Available from: http://www.biotech-asia.org/vol13no2/screening-of-glypican-3-expression-in-human-normal-versus-benign-and-malignant-tissues-a-comparative-study-glypican-3-expression-in-cancers/

3.	Shi D, Shi Y, Kaseb AO, Qi X, Zhang Y, Chi J, et al. Chimeric Antigen Receptor-Glypican-3 T-Cell Therapy for Advanced Hepatocellular Carcinoma: Results of Phase I Trials. Clin Cancer Res [Internet]. Clin Cancer Res; 2020 [cited 2022 Jan 5];26:3979–89. Available from: http://www.ncbi.nlm.nih.gov/pubmed/32371538
