## Supplementary figures and images for "BOXR1030, an anti-GPC3 CAR with exogenous GOT2 expression, shows enhanced T cell metabolism and improved antitumor activity"

### Supplemental Figure S1

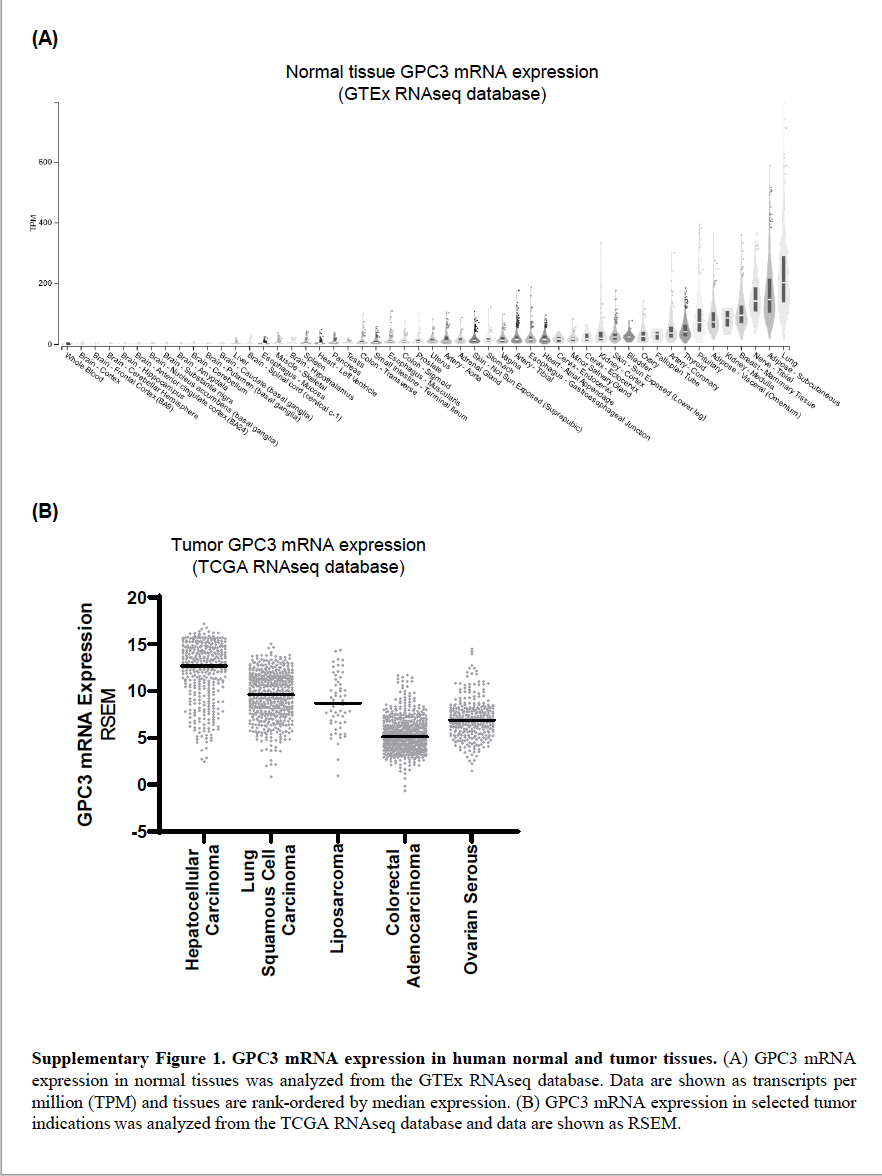

### Supplemental Figure S2

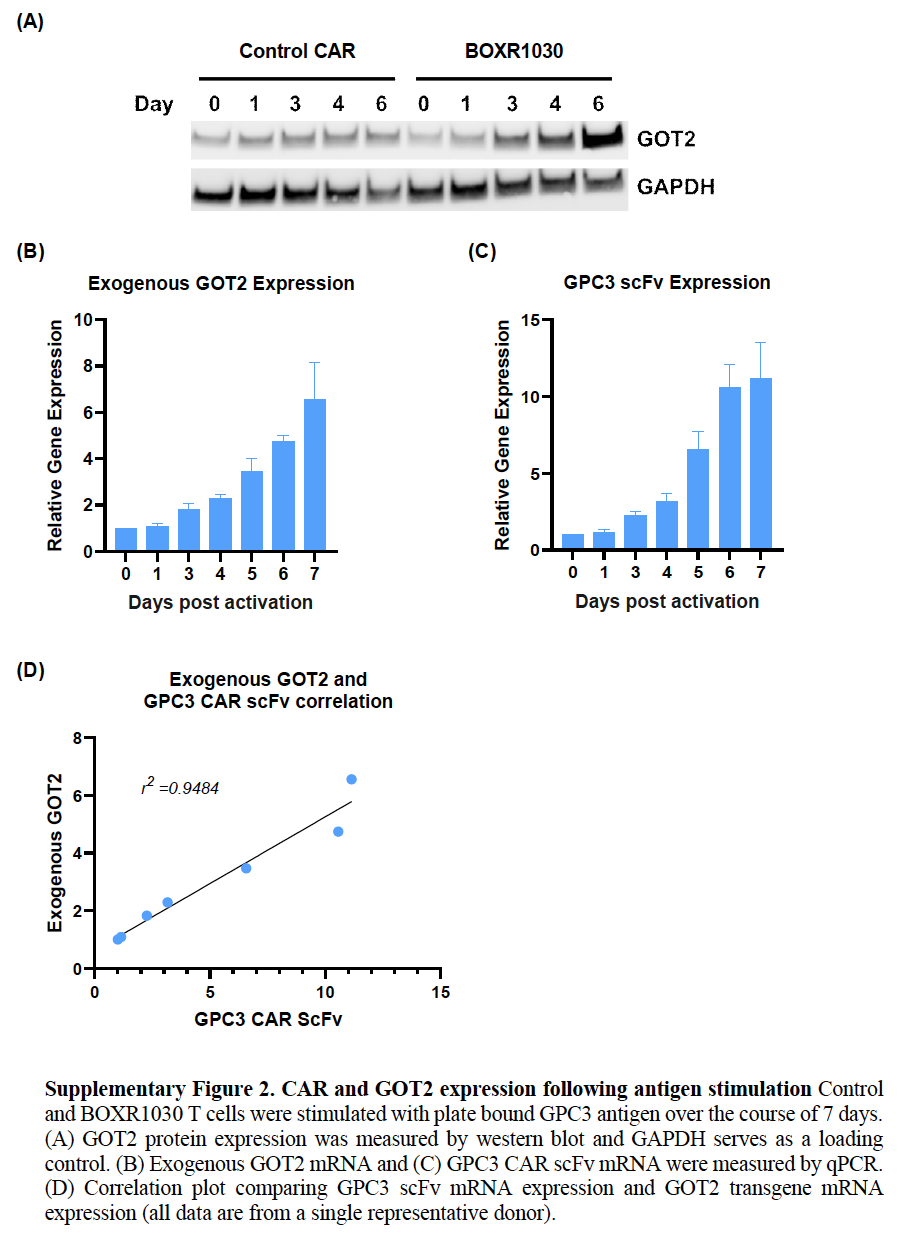

### Supplemental Figure S3

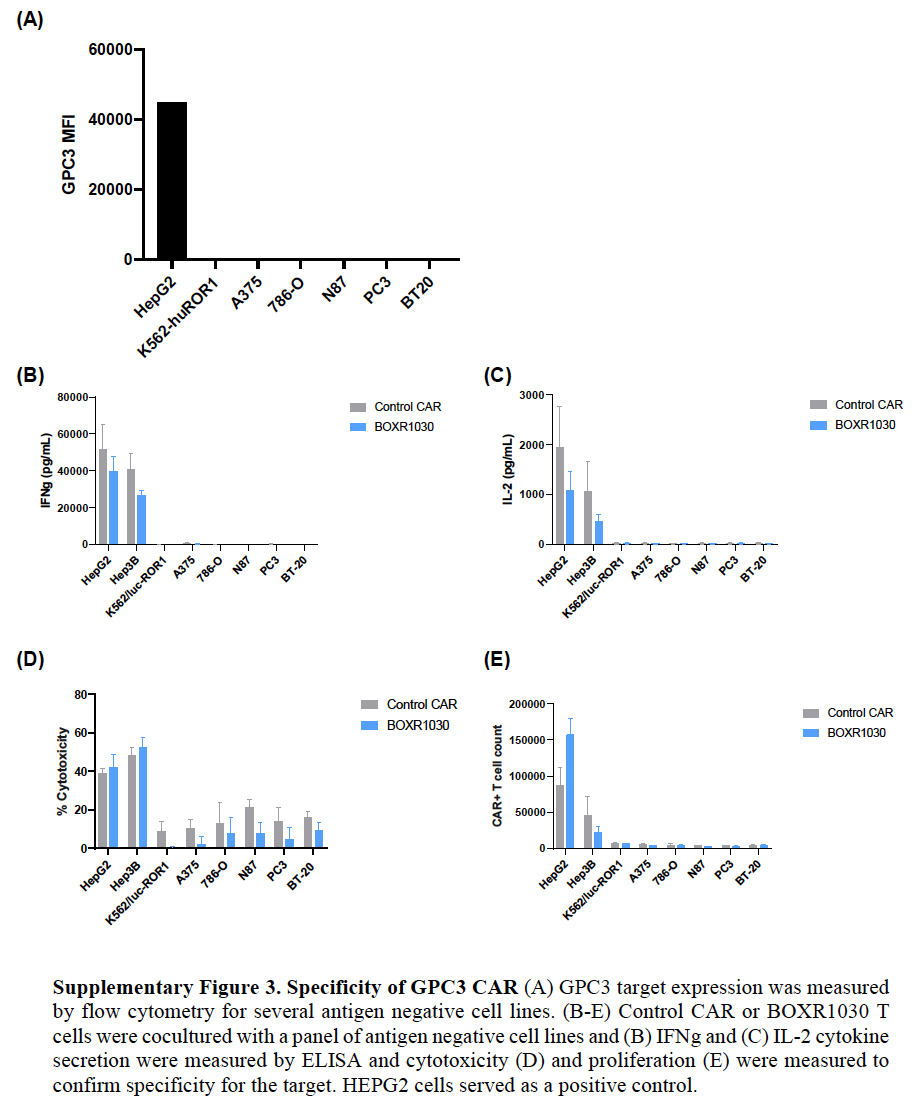

### Supplemental Figure S4

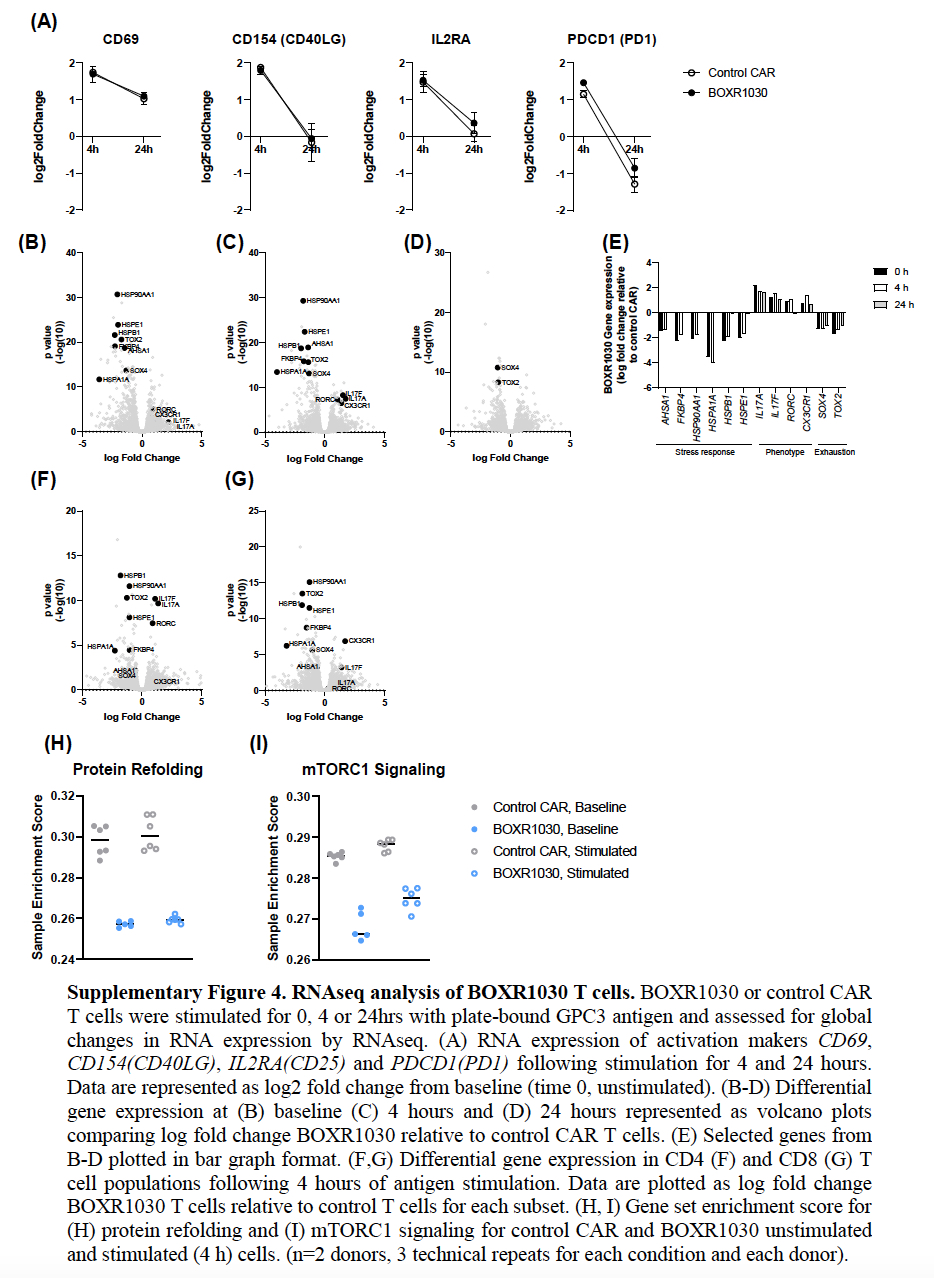
